# Cross-species genomic survey reveals a single origin of the soft-shelled turtles

**DOI:** 10.1101/2024.01.23.576798

**Authors:** Wei Li, Junxian Zhu, Zijian Gao, Liqin Ji, Yanchao Liu, Haiyang Liu, Xiaoli Liu, Xiaoyou Hong, Chengqing Wei, Jiehu Chen, Chen Chen, Xinping Zhu

## Abstract

Turtles have a have a complicated sex determination system; however, Trionychidae species, as a unique group, have evolved a conserved genetic sex determination, which provides an excellent case to study the evolution of environmental sex determination to genetic sex determination. Analysis of the chromosome-level genome assembly, construction of the first sex chromosomes of Trionychidae, and embryonic transcriptome revealed that *P. sinensis* has a ZW-type sex determination system, and that the key sex-determining genes, including *Dmrt1*, *Amh*, *Foxl2*, etc., and their homologs, are not located on the sex chromosomes, which rejects the involvement of *Dmrt1*’s SDGs or dosage compensation effect in the sex determination of *P. sinensis* ZW system and also indicates that the ZW chromosomes in birds, turtles, frogs, and fishes originated from different chromosome pairs in the common ancestors. Subsequently, comparative genomic studies demonstrated that a conserved ZW-type sex determination system exists in Trionychidae species, and inferred that their sex chromosomes originate from the same chromosome pair of the common ancestor. More importantly, we found a single origin of the sex chromosomes in Trionychidae species, approximately 56 million years ago, which is highly coincident with the Paleocene-Eocene thermal maximum, PETM, event. Therefore, we hypothesize that the PETM event and the loss of embryonic self-thermal regulation are two critical dynamics that drive the adaptive evolution of sex chromosomes for the common ancestor of Trionychidae species in response to extreme temperature changes. Finally, we link the ancestral sex chromosome origin time, the species divergence time, the PETM event, and the continental plate movements to paint a picture of a single origin and radiation route for extant Trionychidae species. Collectively, our findings provide novel insights into the mechanisms of sex determination in *P. sinensis*, and the single origin of sex chromosomes and species in extant soft-shelled turtles.

## 1. Introduction

The Chinese soft-shelled turtle (*Pelodiscus sinensis*, Trionychidae), exhibits a ZW-type sex determination system (Kawagoshi et al., 2009; Mu et al., 2015). However, studies on ZW-type sex determination mechanisms have been restricted. Currently, there are two dominant hypotheses. One is the effects of sex-determining genes (SDGs) on the W chromosome. The W-linked DM-domain (*DM-W*) gene was localized on the W chromosome of *Xenopus laevis*, and male tadpoles (ZZ) carried with the *DM-W* expression vectors exhibited ovarian-like gonadal structures (Yoshimoto et al., 2008). The other is the dosage compensation of *Dmrt1* on chromosome Z in birds. Briefly, two doses of *Dmrt1* switched on the male pathway, alternatively, a single dose of *Dmrt1* caused feminization (Ioannidis et al., 2021; Smith et al., 2009). Likewise, *Dmrt1* was first identified as a master gene to be involved in sex determination and/or differentiation in *P. sinensis* (Sun et al., 2017). Meanwhile, the faint evidences of sex chromosome dosage compensation were found in the Trionychidae (Bista et al., 2021; Montiel et al., 2022). However, it remains unclear whether *Dmrt1* is positioned on the sex chromosomes, thereby relying on the dosage compensation (Z chromosome) or SDG (W chromosome) of *Dmrt1* to determine the sex of *P. sinensis*. Because of the lack of detailed genomic information of sex chromosomes, it has raised unprecedented difficulties in the study of the sex determination mechanism of *P. sinensis*.

Moreover, except for *P. sinensis*, previous studies have shown that another species of Trionychidae including, *Apalone spinifera* (Badenhorst et al., 2013), *Rafetus swinhoei* (Ren et al., 2022), and *Chitra indica* (Rovatsos et al., 2017) also possess a ZW genetic sex-determining system, which implies that there is a conservative genetic sex determination system in Trionychidae (Bista and Valenzuela, 2020; Rovatsos *et al*., 2017). This provokes two thoughts. One aspect is how the sex chromosomes originated in Trionychidae and whether they are similar to the XY system in mammals and the ZW system in birds? The classical theory of sex chromosomes evolution suggests that the sex chromosomes originate from a pair autosomes (Bellott et al., 2010; Zhou et al., 2014), which gains sex-determining genes (SDGs) due to mutations or translocations (Bachtrog et al., 2011; Charlesworth et al., 2005). Subsequently, the sexually antagonistic alleles which benefit one sex while damaging the other, such as the genes of controlling male peacock feathers to enhance reproductive advantages but making females more susceptible to capture (Zhu et al., 2022), gradually accumulate around them thereby exacerbating recombination suppression in favor of linkage heredity of these genes with SDGs in one sex (Rice, 1987). This process results in the degradation and loss of functions of a large number of genes on Y or W, thereby forming heterochromatin (Bachtrog et al., 2014; Wright et al., 2016). Most of the Y and W regions have stopped recombining with the X and Z chromosomes; however, some recombination regions (called the pseudoautosomal region) which are usually located at the ends of the chromosomes, remain in order to allow normal pairing during meiosis (Vicoso, 2019; Xu et al., 2019b).

On the other hand, the soft-shelled turtles with a conserved ZW sex-determining system are widely distributed all over the world, such as *P. sinensis* mainly in China, *A. spinifera* largely in the United States, *R. swinhoei* primarily in China and Vietnam, and *C. indica* mostly in Bangladesh, India, Nepal, and Pakistan (Shine, 2013). Additionally, the soft-shelled turtles are also spread in parts of Africa. Intriguingly, there is no distribution of the soft-shelled turtles in South America and Europe at the same latitude (Shine, 2013). Whether turtles with the ZW sex-determination system acquired their sex chromosomes in their common ancestors and then dispersed everywhere; alternatively, turtles everywhere experienced parallel evolution (Bolnick et al., 2018; Cheng et al., 2021) of the ZW sex chromosome system under similar selective pressures, after spreading out from their common ancestors?

All extant turtles share the common ancestors (Li et al., 2018; Schoch and Sues, 2015). The earliest ancestors with relatively complete soft-shell and hard-shell turtle characteristics can be traced back to ∼120 million (Li et al., 2015) and ∼210 million years ago (Shine, 2013), respectively, suggesting that turtles finished their major evolutionary processes in the hundreds of millions of years ago. However, turtles display complex and diverse sex determination systems (Johnson Pokorná and Kratochvíl, 2016; Thépot, 2021). Environmental sex determination (ESD) is prevalent in most turtles, and the predominant environmental factor is temperature, named temperature-dependent sex determination (TSD) (Li and Gui, 2018). By contrast, part of turtles possesses genetic sex determination (GSD), such as *A. spinifera*, *P. sinensis*, and *R. swinhoei* of Trionychidae hold a ZW microchromosome system (Badenhorst *et al*., 2013; Mu *et al*., 2015; Ren *et al*., 2022). More importantly, different sex determination patterns (TSD, *Carettochelys insculpta* and GSD) co-exist in Trionychia (Webb et al., 1986). Compared to the conserved sex determination systems in mammals (XY) and birds (ZW), turtles present multiple sex determination methods and are considered to be a special group that has transitioned from ESD to GSD (Capel, 2017; Sarre et al., 2011). Consequently, turtles are a suitable model for studying sex chromosomes origin and evolution.

In this study, we construct a chromosome-level genome of *P. sinensis* and then synthesized four soft-shelled turtles’ genome resequencing and *P. sinensis* embryonic transcriptome data to clarify the first sex chromosomes in Trionychidae. Our study provides a novel insight into the sex-determination mechanism of *P. sinensis* and the fundamental data on the origin and evolution of sex chromosomes in Trionychidae, helping to unravel the mystery of the evolution from ESD to GSD in Testudines. More importantly, we also provide a clue to the single origin for the soft-shelled turtle.

## 2. Materials and Methods

### 2.1 Sample collection, library construction and sequencing

For whole genome sequencing, genomic DNAs were extracted from the muscle tissues from a mature female *P. sinensis*, cultivated in Guangzhou, China. To perform the genome shotgun sequencing, a 350-bp insert library was constructed, followed by paired-end sequencing (2 × 150 bp) on an Illumina Novaseq 6000 platform (Illumina, San Diego, CA, USA). For the long-read sequencing, a Nanopore library with an insert-size of 40 kb was prepared, and subsequently sequenced on a Nanopore platform (Nanopore Technologies, Oxford, Head Office, UK). The Hi-C library, constructed with the same DNA sample, was sequenced on the same Illumina Novaseq 6000 platform.

In order to assist genome annotation, total RNAs were extracted from various tissues of the female *P. sinensis*, including muscle, brain, kidney, spleen, liver, and gonads. Individual cDNA library was constructed for transcriptome sequencing (RNA-seq) on the Pacbio platform (Pacific Biosciences, Menlo Park, CA, USA). In addition, to explore gene expression differences during sex development, three male and three female samples were collected from *P. sinensis* embryo at each embryonic stage. In brief, the samples were stripped away from the eggshells on days 3, 4, 5, 6, 7, 8, 9, 13, 14, 15, 16, 18, 22, 26 and 37, and sex of the embryos was identified via PCR amplification after the DNA extraction. RNAs from each sample were extracted, followed by cDNA library construction for transcriptome sequencing (RNA-seq) on the Novaseq 6000 platform.

Moreover, for the genome-wide association analysis, 15 males and 15 females from each species, including *P. sinensi*s, *Palea steindachneri*, *Apalone spinifera*, and *Apalone ferox* were collected. The genomic DNAs were also extracted from the muscle tissues to construct 500-bp insert libraries, which were then sequenced on Illumina Novaseq 6000 platform.

### 2.2 Genome size estimation based on the routine k-mer method

For filtering of short reads, reads with N content over 10% or low quality (Q<=5) base over 50% are considered as low-quality reads, and corresponding paried-end reads will be discarded. Cleaned short reads were then used for genome size estimation based on a 21-mer distribution with jellyfish. The genome size, heterozygosity, and repetitiveness were estimated using GenomeScope 2.0.

### 2.3 Genome assembly and chromosome construction

For the genome assembly, Canu (v2.2, -nanopore, genomeSize = 2 G) was employed to assemble the long reads to super contigs. To construct the chromosomes, those low mapping quality reads, multiple hits and singletons of Hi-C data were discarded with HiCUP. Subsequently, clean reads were mapped to the super contigs via BWA (v0.7.17-r1188). Mapped reads were further clustered using Juicer, followed by ordering and orientation with 3d-dna to obtain the chromosome-level genome. By Hic-Pro (v3.1.0), the Hi-C data was aligned with the genome to construct an interaction matrix, which was then used to generate a contig interaction matrix figure with Hicplotter (v0.6.6). For genome polishing, Pilon (v1.24) was applied for calibration of the genome along with the cleaned short reads.

To assess the quality of our final chromosome-level genome assembly, contig N50 and scaffold N50 values were calculated, and with the popular sauropsida_odb10 database, BUSCO (v5.2.2) was employed to evaluate the completeness of the assembled *P. sinensis* genome.

### 2.4 Genome annotations and gene predictions

Prediction of repeat elements was based on homology methods using RepeatMasker. The rmblastn search engine was employed to search the repeat element with default parameters. The distribution of repetitive sequences is calculated and plotted using an internal python script with a 50 Kb statistical window and 25 Kb steps.

Genes were annotated on the repeat masked genome using the MAKER genome annotation pipeline (v. 3.01.02), by three approaches, de novo, homology and transcriptome-based. For the de novo prediction, SNAP and Augustus were applied with default parameters. For the homology-based prediction, protein sequences of 11 representative species, including *Caretta caretta*, *Chelonia mydas*, *Dermochelys coriacea*, *Chelydra serpentina*, *Chrysemys picta*, *Terrapene carolina triunguis*, *Trachemys scripta elegans*, *Mauremys mutica*, *Mauremys reevesii*, *Platysternon megacephalum*, *Chelonoidis abingdonii* and *Gopherus evgoodei* were downloaded from Ensembl for mapping to our assembled genome using TBLASTn. Subsequently, GeneWise was used to predict gene structures of the achieved alignments. For the transcriptome-based prediction, ccs 4.0.0 (https://github.com/PacificBiosciences/ccs) was used to generate the circular consensus sequences with the subreads. After classification and clustering by lima (v1.10.0, https://github.com/pacificbiosciences/barcoding/) and isoseq (v3, https://github.com/PacificBiosciences/IsoSeq), the high-quality polished consensus isoforms were aligned with the genome by using minimap2 to obtain the predicted genes. Finally, these predicted results were integrated using MAKER to obtain a final consistent gene set.

To perform functional annotation, BLASTp was applied to align the predicted protein sequences against four public databases, including SwissProt, TrEMBL, KEGG and InterPro. These results were retrieved using Gene Ontology (GO) terms.

### 2.5 Variation detection and genome-wide association analysis

The whole genome re-sequencing reads from 4 species were mapped to the assembled *P. sinensis* genome by BWA (0.7.17-r1188). Samtools (v1.14) was applied to estimate the mapping depth, identity for each species. Variant calling was analyzed via deepvariant (v1.40) with WGS model. The generated VCF file was then converted to a Ped/Map format using PLINK software, which was also used to filter out individuals with an individual variant missing rate (-mind) > 10%, a missing rate per SNP (-geno) > 10%, minimum allele frequency (-maf) < 5%.

As the phenotypic is a binary trait (male and female were recorded as 0 and 1, respectively), genome-wide association study of the phenotypic and SNP data was performed using general linear model (GLM) with the R package GAPIT3 (v3.3). LD blocks analysis was performed via LDBlockShow with default parameters.

### 2.6 Evolution analysis

To construct orthogroups from protein-coding genes of *P. sinensis*, we downloaded CDS of several representative species from Genbank. After multiple sequence alignment with predicted CDS of *P. sinensis* and other species using BLASTp (e-value ≤ 1e-5), orthogroups were clustered with OrthoFinder (v2.5.4). In order to reveal the phylogenetic position of *P. sinensis*, we employed mafft (v7.490, maximum number of iterative refinement = 1000) with local-pair method to align the CDS sequences of single-copy orthogroups.

After obtaining the conserved regions via trimAl (v1.4. rev15), a phylogenetic tree was constructed using IQtree (v2.0.5) with the aligned CDS of all single-copy genes, based on the best model for each gene selected by ModelFinder tool without subsequent tree. Subsequently, fossil-calibrated nodes from TIMETREE in the phylogenetic topology, including 247.1–260.2 Mya between Alligator sinensis and Gallus gallus were set. based on all exons over 500 bp from single-copy orthogroups, the BEAST (v2.7.1) was applied with HKY+G model and 10,000,000 iterations to estimate the divergence time.

Chromosome colinearity analysis was performed using JCVI utility libraries (version: v1.2.20, https://github.com/tanghaibao/jcvi) (The JCVI software package is the Python version of the McScan software.). Based on the coding gene sequence, the ortholog method of JCVI was used to establish the corresponding relationship between genes in the genome. The alignment of homologous genes was performed using Lastal, with c-score cutoff of 0.7, and the minimum number of anchors in a cluster of 4. Homologous genes were selected, and genes with blocks with anchors >= 40 were selected for chromosome colinearity mapping.

### 2.7 Estimation for the formation time of the sex chromosomes

Assuming that the chromosomes of these species evolved independently at a rate equal to the lineage-specific rate of divergence, the formation times of specific chromosomes for each species can be estimated based on the Ks method. With the CDS of the single-copy orthogroups, seqinr (v4.2-30) was used to calculate the Ks for single copy genes between *P. sinensis* and other species. The mean lineage specific divergence rate of *P. sinensis* lineage was calculated according to the following formula: Mean Lineage Specific Divergence Rate = Lineage-specific Median Ks / Corresponding Divergence Time of Species. The formation time of the sex chromosomes was estimated as the product of the median Ks for single-copy genes in sex chromosomes and the mean lineage specific divergence rate.

### 2.8 Differential gene expression analysis

Comparative quantitative gene expression analysis was performed using RSEM, and gene expression was calculated using the fragments per kilobase per million reads (FPKM) method, where FPKM=cDNA fragments/mapped fragments (millions)/transcript length (kb). Then, the sequencing results (each group included three biological replicates) were analysed using DESeq2 R (1.34.0). In the DEG assay, the screening criteria were a fold change ≥ 1.5 and FDR < 0.05. The functional annotation and classification of DEGs were conducted with the GO database and KEGG database and via the analysis of DEG enrichment metabolic pathways. Protein-protein interaction networks (PPI) analysed using STRING (v11.5, https://string-db.org), with Pelodiscus sinensis database and minimum required interaction was 0.400 (medium confidence), then analyzed by MCODE. Functional enrichments about sex differentiation in the network was selected with false discovery rate (FDR) < 0.05 on top 10 strength with gene ontology, KEGG pathways and local network cluster (STRING). Cytoscape (http://cytoscape.org/) was used for visualization the network.

## 3. Results

### 3.1 High-quality genome assembly and annotation

In total, 121.67 Gb of female and 136.56 Gb of male clean data from whole-genome sequencing were used to assess the genome size, heterozygosity, and repeat proportion of *P. sinensis*. Based on the distribution of 21-mers in female and male, the dominant peak depth was 40 and 47, the estimated genome size was approximately 1.95 Gb and 1.85 Gb, the heterozygosity was ∼0.70% and ∼0.88%, and the repeat proportion was ∼20.12% and ∼18.21%, respectively. From the Oxford Nanopore sequencing of female, we collectively obtained 137.24 Gb of raw data with an average subreads length and N50 length of 22,275 bp and 31,520 bp, respectively. The Oxford Nanopore long consecutive reads were used for de novo assembly and the Illumina short reads were applied for polishing, then we obtained an initially assembled genome (contig) with a size of 2.20 Gb consisted of 19,904 contigs, a contig N50 of 3.36 Mb, and a GC content of 44.41%. Additionally, we constructed two Hi-C libraries and gained a total of 141.81 and 140.55 Gb clean data to anchor the assembly contigs into chromosomal-scale genome, with a genome size of 2.36 Gb, a N50 of 124.95 Mb, and a GC content of 42.74%. Final assembled genome contained 32 autosomes and the sex chromosomes of Z and W, of which the chromosome length ranged from 1.38 Mb (Chromosome 32, Chr32) to 334.17 Mb (Chromosome 1, Chr1) (Figure 1). Illumina short reads from female and male libraries were aligned back to the assembled *P. sinensis* genome, and the alignment rates were 98.39% and 98.43%, respectively. The genome completeness was evaluated using BUSCO and the results revealed that the assembly genome consisted of 7,112 conserved core genes (95.08), including 94.17% of complete and single-copy BUSCOs (S), 0.91% complete and duplicated BUSCOs (D), 1.30% Fragmented BUSCOs (F), and 3.62% Missing BUSCOs (M).

**Figure 1.**
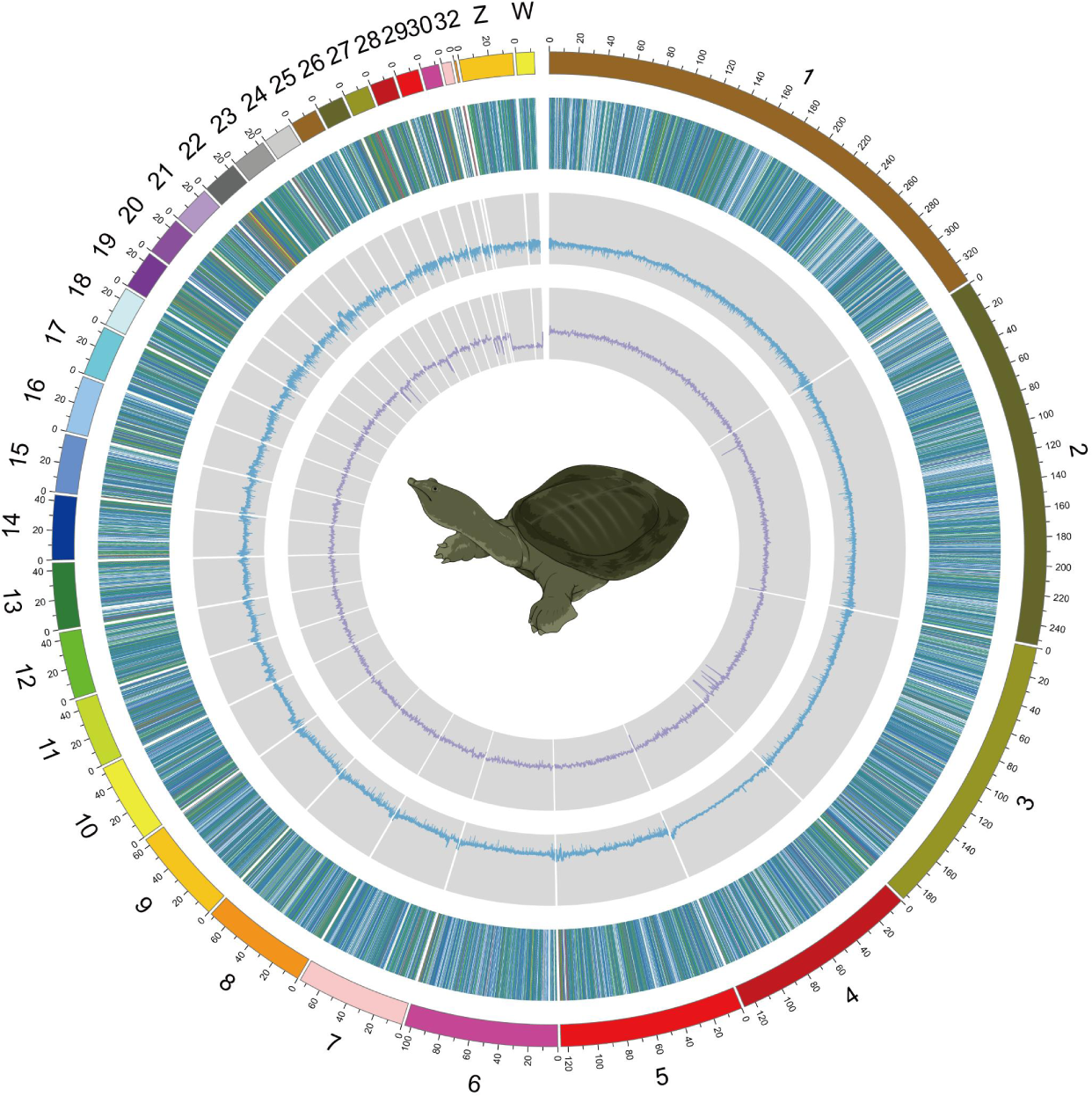
Genome assembly and annotation of *P. sinensis*.

A total of 1,056,413,370 bp of repetitive sequences were identified, accounting for 44.83% of the assembled genome size using the RepeatMasker database (version 3.7). The percentages of DNA transposons and retroelements in the genome was 23.96% and 17.68%, respectively, of which the long interspersed nuclear elements (LINEs) were the most abundant DNA transposons of 10.25%, followed by the long terminal repeats (LTRs) of 6.44%, and the short interspersed repetitive elements (SINEs) of 0.99%. Simultaneously, a number of unclassified repetitive sequences of 1.77% were found in the genome. Moreover, we identified a total of 30,829 protein-coding genes in *P. sinensis* genome, and the average lengths of coding genes, introns, exons, coding sequences (CDSs) were 21.88 kb, 7.87 kb, 288 bp, and 164 bp, respectively. Among these genes, 13,468 genes were functionally annotated in GO, KEGG, KOG, Uniport, and NR databases. Meanwhile, many non-protein coding genes were also identified, including 5 rRNA, 509 tRNA, and 942 microRNA genes.

### 3.2 Resequencing of four typical species in Trionychidae

A total of 30 *P. sinensis*, 30 *P. steindachneri*, 30 *A. spinifera*, and 30 *A. ferox* were collected for resequencing analysis, then the resequencing data of four soft-shelled turtles were mapped on the *P. sinensis* genome and the results presented that the depth of *P. steindachneri*, *A. spinifera*, and *A. ferox* ranged from 6.92 to 13.26, 6.79 to 16.55, and 7.02 to 15.96, respectively (Figure 2A). The identity of *P. steindachneri*, *A. spinifera*, and *A. ferox* with *P. sinensis* genome ranged from 93.94 to 95.36%, 92.23 to 94.42%, and 92.17 to 94.54%, respectively (Figure 2B). The coverage of *P. steindachneri*, *A. spinifera*, and *A. ferox* with *P. sinensis* genome ranged from 60.00 to 89.76%, 56.57 to 86.95%, and 56.81 to 86.59%, respectively (Figure 2C). Remarkably, in the three soft-shelled turtles, both female and male genomic data detected highly homologous sequences to the Z and W chromosomes of *P. sinensis*, and the depth of W chromosome in females was approximately half that of the Z chromosome in males. Additionally, the results also found that 26,923 SNPs in *P. sinensis* (Figure 2D), 25,817 in *P. steindachneri* (Figure 2E), 31,294 in *A. spinifera* (Figure 2F), and 14,542 in *A. ferox* (Figure 2G) showed significant association with sex and all SNPs were localized to sex chromosomes, all showing heterozygosity in females and purity in males. Taken together, the above results implied that *P. steindachneri*, *A. spinifera*, and *A. ferox* had a sex determination system of female heterogamy as similar to that of *P. sinensis*.

**Figure 2.**
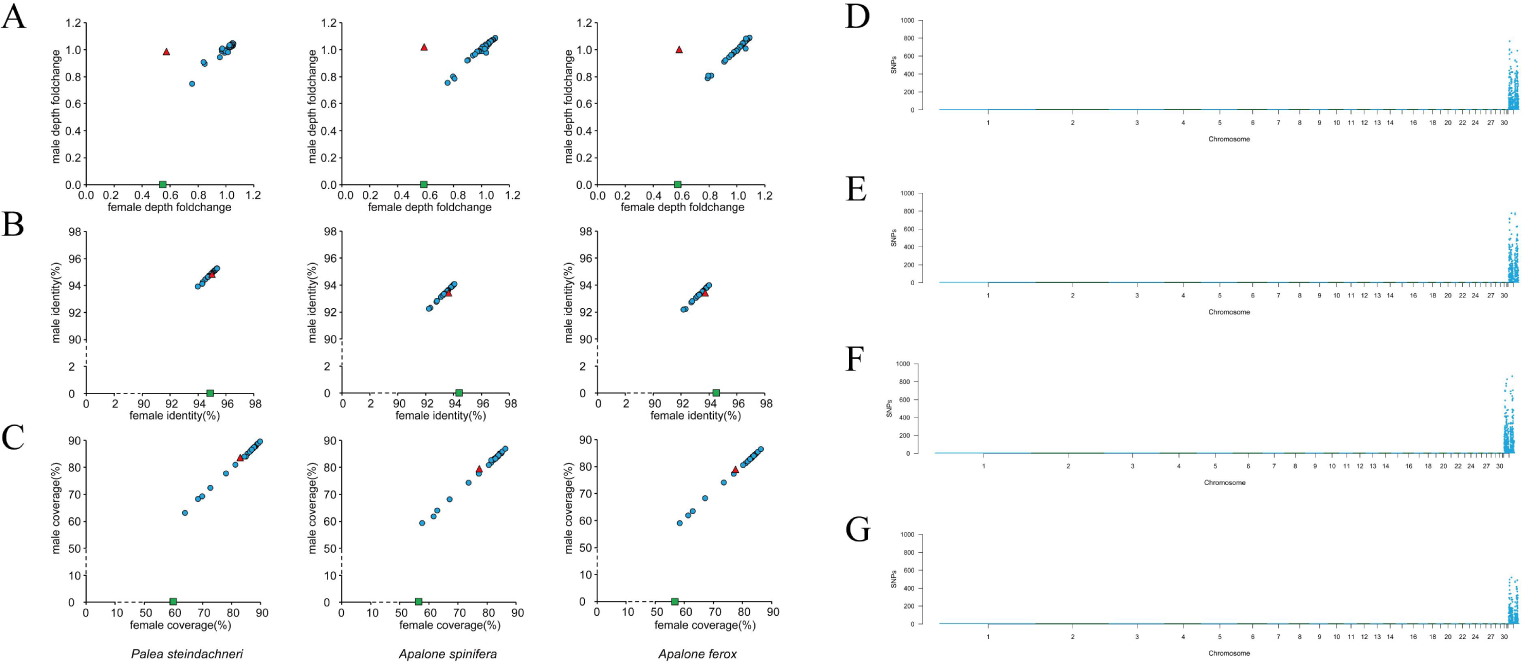
Whole-genome sequencing to analyze the sequencing depth (A), genome similarity (B), and coverage (C) of *P. steindachneri*, *A. spinifera*, and *A. ferox* to *P. sinensis*, and the density of sex-specific SNPs in *P. sinensis* (D), *P. steindachneri* (E), *A. spinifera* (F), and *A. ferox* (G).

### 3.3 Phylogenetic relationship and divergence time evaluation

We used a total of 6713 single-copy orthologs from *P. sinensis* to construct a ML phylogenetic tree with 29 other turtles, and *Alligator sinensis* and *Gallus gallus* as outgroups. The results showed that *P. sinensis* was most closely related *Palea steindachneri*, followed by *Rafetus swinhoei*, *Apalone spinifera*, *Apalone ferox*, and *Pelochelys cantorii*, then clustered within the family of Trionychia, together with *Carettochelys insculpta* (Figure 3A). Subsequently, A molecular clock was built to estimate the divergence time according to the present fossil record, showing that the ancestor of *P. sinensis* separated from the ancestor of *P. steindachneri* approximately 13.16∼13.68 million years ago. The ancestor of *P. sinensis*, *P. steindachneri*, *R. swinhoei*, *A. spinifera*, *A. ferox*, and *P. cantorii* separated from the ancestor of *Carettochelys insculpta* approximately 69.19∼71.53 million years ago. Moreover, the ancestor of the GSD turtles (or soft-shelled turtles) separated from the ancestor of the TSD turtles (hard-shelled turtles) approximately 80.71∼82.88 million years ago (Figure 3B).

**Figure 3.**
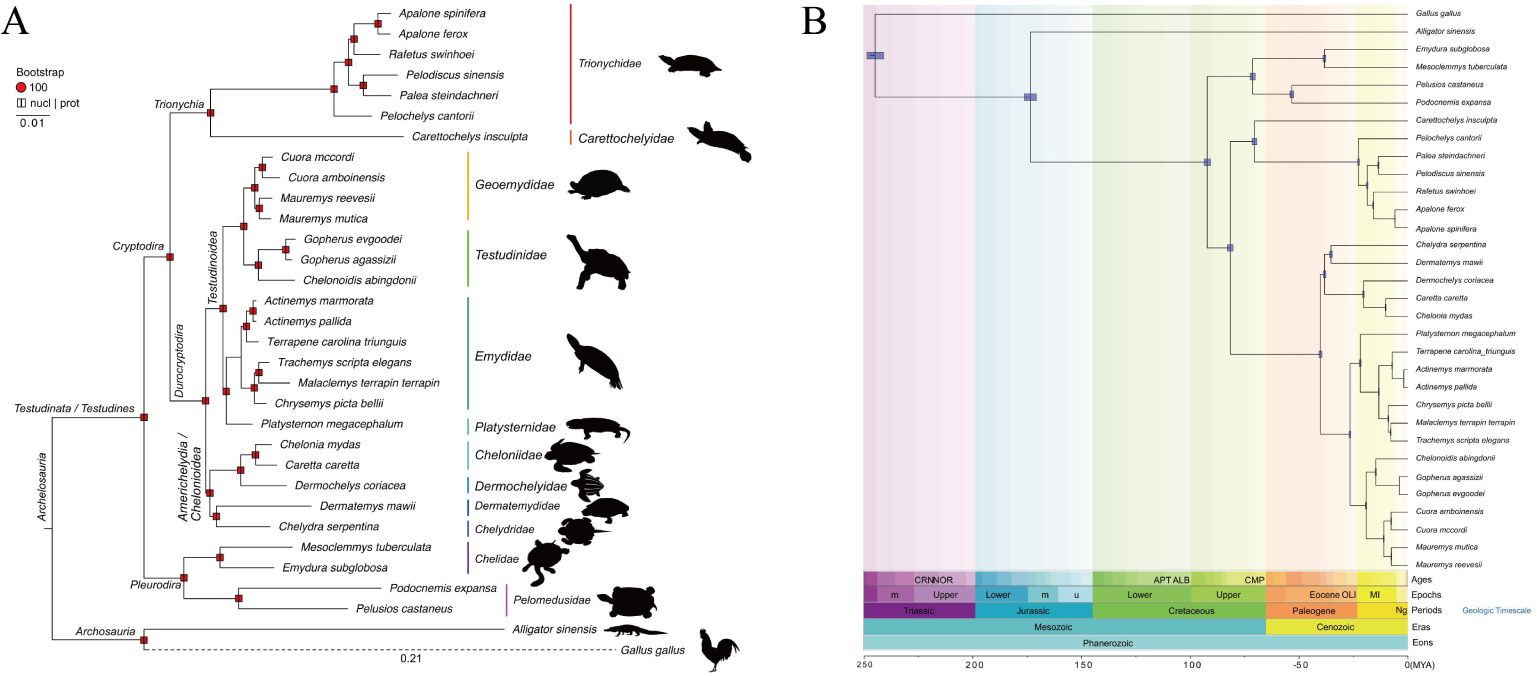
Phylogenetic relationships (A) and species divergence time in turtles (B)

### 3.4 Origin time of the sex chromosomes

Based on the genome-wide coding sequences, the lineage-specific median Ks between *P. sinensis* and the nine hard-shelled turtles, including *T. carolina triunguis*, *M. mutica*, *C. mydas*, *C. picta bellii*, *M. reevesii*, *T. scripta elegans*, *G. evgoodei*, *C. abingdonii*, and *D. coriacea*, was estimated to be 0.2031 and the divergence time between *P. sinensis* and the nine hard-shelled turtles ranged from ∼80.17 to 82.88 million years ago (Mya). Therefore, the mean lineage-specific divergence rate for the *P. sinensis* could be calculated to be between 0.00245 and 0.00253 change/site/Mya according to this information. Moreover, the mean value of Ks between the Z and W sex chromosomes of *P. sinensis* was 0.2616. Assuming that the Z and W sex chromosomes of *P. sinensis* evolved independently at a rate equal to the lineage-specific rate of divergence (Pan et al., 2019), the combined divergence rate would be between 0.0049 and 0.0051 change/site/Mya. Eventually, we obtained the approximate differentiation time of the Z and W sex chromosomes in *P. sinensis* between 56 and 48 Mya (Figure 4).

**Figure 4.**
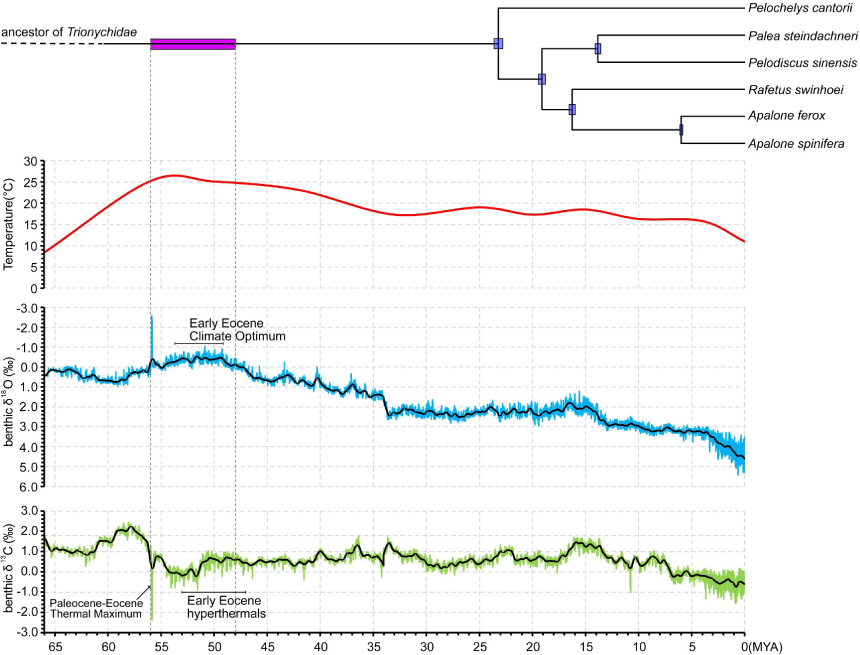
Estimating the time of the sex chromosome origin in *P. sinensis*.

### 3.5 Evolution of the sex chromosomes

To investigate the origin of *P. sinensis* sex chromosomes, the synteny analysis was processed between the GSD and TSD turtle genomes. Concretely, *P. sinensis* genome presented a high degree of collinearity with the genomes of other turtles, including *Mauremys reevesii*, *Mauremys mutica*, *Gopherus evgoodei*, *Trachemys scripta elegans*, *Chelonia mydas*, *Caretta caretta*, and *Dermochelys coriacea*. Additionally, Chromosome inversions, deletions, rearrangement, and fission events were observed during the evolution from TSD (or hard-shelled turtles) to GSD (or soft-shelled turtles) turtles. Significantly, the Z and W chromosomes of *P. sinensis* derived from the chromosome 18 Chr of *Mauremys reevesii*, 16 Chr of *Mauremys mutica*, 13 Chr of *Gopherus evgoodei*, 15 Chr of *Trachemys scripta elegans*, *Chelonia mydas*, *Caretta caretta*, and *Dermochelys coriacea*. Furthermore, these chromosomes were highly conserved during evolution from TSD (or hard-shelled turtles) to GSD (or soft-shelled turtles) turtles. More importantly, we found that the key genes involved in sex determination of *P. sinensis* including *Dmrt1*, *Foxl2*, *Rspo1*, *Sox8*, *Esr1*, and *Amh*, were not distributed on the sex chromosomes (Figure 5A). Compared with the original sex chromosomes of Testudines, the primary characteristics of *P. sinensis* sex chromosomes were the presence of two obvious inversions at both ends of the Z chromosome, and the loss of coding genes and shortening in the W chromosome (Figure 5B). Meanwhile, the analysis of nonsynonymous mutation frequency (Ka) and synonymous mutation frequency (Ks) of chromosome-encoded genes showed that the softshell turtle W chromosome had the highest values of Ka/Ks compared to autosomes, followed by Z chromosome (Figure 5C and D). An investigation of the distribution of repetitive elements in *P. sinensis* chromosomes revealed that the LINE element was significantly accumulated at one end of the W chromosome (Figure 5E).

**Figure 5.**
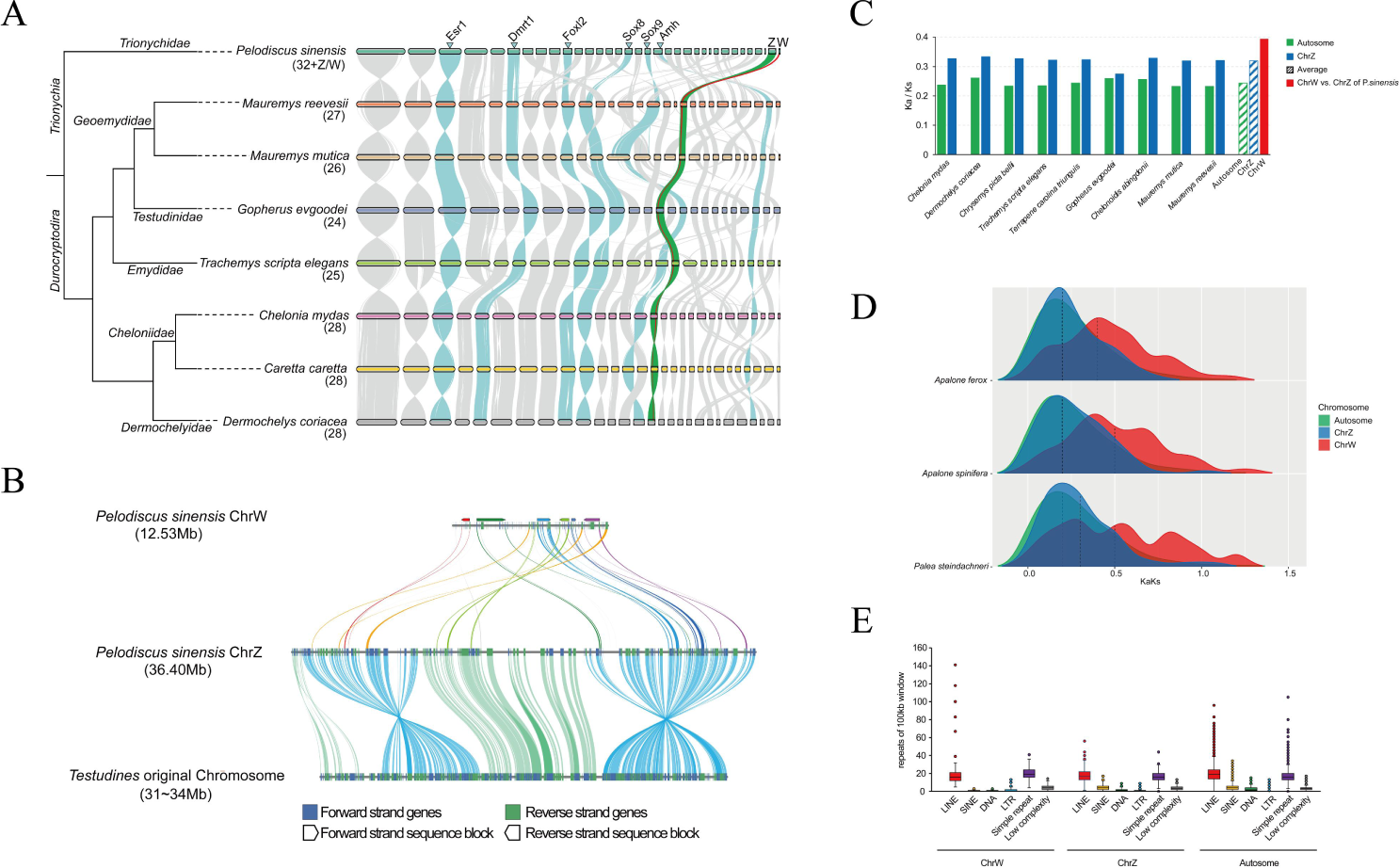
Evolution of the sex chromosomes in *P. sinensis*. Plate A and B represent the synteny analysis of the genomes between *P. sinensis* and seven other turtles, and between *P. sinensis* sex chromosomes and the original sex chromosomes of Testudines, respectively. The values of Ka/Ks for the chromosomes of nine turtles and *P. sinensis*, and three soft-shelled turtles are shown in plate C and D, respectively. Plate D indicates the distribution of repetitive elements in *P. sinensis* genome.

### 3.6 Transcriptome landscape during embryonic development of *P. sinensis*

Gene expression pattern is intimately linked to gene functions; therefore, we performed a transcriptome analysis of the female and male gonads during the embryonic development to uncover the differential expression genes of *P. sinensis* across the sex differentiation process. Heatmap results revealed that genes of the W chromosome displayed differential expression before the putative critical period of sex determination, earlier than the sex determination master genes which were identified in previous studies in *P. sinensis*, including *Dmrt1*, *Amh*, and *Foxl2*. Conversely, most genes on the Z chromosome have shown differential expression after the putative critical time of sex determination (Figure 6A). WGCNA analysis displayed that the tan block (dominated by W chromosome genes) was significantly correlated with the female sex of *P. sinensis* with the highest positive correlation, while the purple block (dominated by Z chromosome genes) had the highest negative correlation. Additionally, the key genes (autosomal genes) for *P. sinensis* sex determination identified in previous studies, *Dmrt1* and *Amh* were located in the midnightblue block (Figure 6B). The results of the protein interaction network demonstrated that there were interactions among W chromosome genes, Z chromosome genes, and autosomal genes (sex-determination key genes) (Figure 6C), and that the trends of their differential expression responded to time; briefly, the W genes were the first to be differentially expressed, followed by Z chromosome genes, and finally to the block of sex-determination key genes (Figure 6D). Collectively. the findings suggested that there might be sex-determining genes on chromosome W that regulated *Dmrt1* or *Amh*, thus allowing the bipotential gonads to differentiate into testis and ovaries.

**Figure 6.**
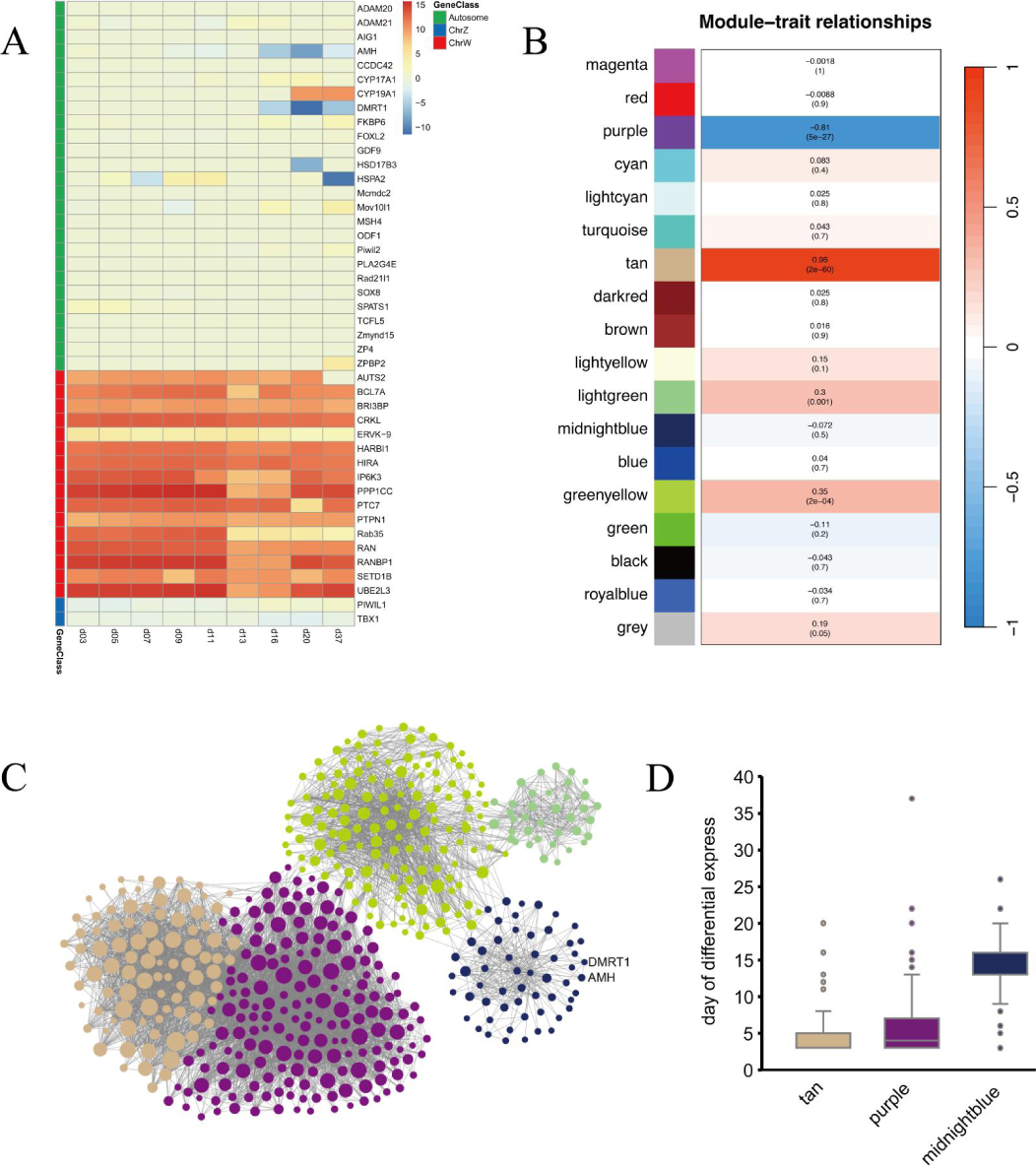
Transcriptome analysis of the expression of genes related to sex determination during embryonic development of *P. sinensis*. Plate A represents a heatmap of differential expression genes from autosomes, Z, and W chromosomes. WGCNA analysis of expression blocks significantly associated with female sex is shown in plate B. Plate C indicates the protein interactions between W, Z, and autosomal genes. Plate D shows the time tendency of differentially expressed genes in the W, Z, and midnightblue block.

## 4. Discussion

### 4.1 The first sex chromosomes of Trionychidae

Assembly and annotation of high-quality and complete genome are the bases for biological researches, bringing a novel dawn to the investigation of sex determination mechanisms and vertebrate studies moves into the genomic era with the opening of the Vertebrate Genome Project (Rhie et al., 2021). In *Scophthalmus maximus*, a chromosomal-level genomic association analysis identified *Sox2* as a major driver in the ZZ/ZW-type sex determination system (Martínez et al., 2021). Chromosome-level genome assembly of *Cynoglossus semilaevis* confirmed the male sex determination gene *Dmrt1* on chromosome Z, and regulated the sex determination of *C. semilaevis* via the dosage compensation of *Dmrt1* (Chen et al., 2014). Also, genome sequencing of XX, XY, and YY southern catfish (*Silurus meridionalis*) revealed that the male sex-determining gene *Amhr2y* was distributed on chromosome 24 and knockout of *Amhr2y* resulted in male to female sex reversal (Zheng et al., 2022). However, studies on the genome of *P*. *sinensis* have been stagnant at the draft genome level for a long time (Wang et al., 2013). In this study, we constructed the first chromosome-level genome of *P*. *sinensis*, combining Oxford Nanopore sequencing, Illumina, and Hi-C technologies, with the genome size of 2.36 Gb and the scaffold N50 124.95 Mb, which is larger than the reported Trionychidae species *P*. *sinensis* (Wang *et al*., 2013), *A. spinifera* (Gessler et al., 2023), *R. swinhoei* (Ren *et al*., 2022), and *P. cantorii* (Liu et al., 2023) genomes and even other hard-shelled turtles including *Trachemys scripta* (Brian Simison et al., 2020), *Chelonia mydas*, and *Dermochelys coriacea*, providing an invaluable genetic resource for the origin and evolution of Trionychidae. The assembled *P*. *sinensis* genome has 32 autosomes, and complete Z and W sex chromosomes, which directly demonstrated that *P*. *sinensis* turtle has a ZW-type GSD system, matching with previous studies (Liang et al., 2019; Mu *et al*., 2015). Additionally, it also signifies that we have constructed the first sex chromosomes in Trionychidae. Although the chromosome-level genome was also possessed in *R. swinhoei* and *P. cantorii*, the complete sex chromosomes have not been assembled (Liu *et al*., 2023; Ren *et al*., 2022). Moreover, *P*. *sinensis* and *A. spinifera* currently only have draft genomes (Gessler *et al*., 2023; Wang *et al*., 2013).

Mechanisms of sex determination in turtles are complex and diverse, and undergo frequent and independent evolution, varying even among closely related species (Bista and Valenzuela, 2020; Thépot, 2021), with the Trionychidae species providing a prime example for study. *Dmrt1* was the first major gene identified to regulate sex determination in *P*. *sinensis*, which was differentially expressed during the critical periods of sex determination, and knockdown of *Dmrt1* resulted in male to female sex reversal (Sun *et al*., 2017). Similarly in most vertebrates, *Dmrt1* and its homologs have been characterized as a key player in sex determination (Ge et al., 2018; Ioannidis *et al*., 2021; Kondo et al., 2003; Matson et al., 2011). *Dmrt1* and its homologs are involved in sex determination in the ZW-type sex determination system mainly through the effects of the SDGs (Yoshimoto *et al*., 2008) or the dosage compensation (Chen *et al*., 2014; Smith *et al*., 2009). However, our findings showed that *Dmrt1* (or its homologs) was not located on the sex chromosomes. Moreover, the results of the WGCNA revealed that the genes on the W chromosome were significantly correlated with the sex of *P*. *sinensis*, while the genes on the Z chromosome were strongly negatively correlated. Meanwhile, the W chromosome genes were the first to undergo differential expression in response to sex differentiation, followed by Z chromosome genes, and finally to the block of *Dmrt1* gene. Therefore, we rejected the hypothesis that *Dmrt1* (or its homologs) participates in the sex determination of *P*. *sinensis* via its SDGs or dosage compensation effects. The above results suggest that the ZW chromosomes in birds, turtles, frogs, and fishes originated from different chromosome pairs in the common ancestors and may have undergone multiple independent origins (Matsubara et al., 2006; Vallender and Lahn, 2006). We also found the similar patterns in other star genes for sex determination in *P*. *sinensis*, such as *Amh* (Zhou et al., 2019), *Foxl2* (Jin et al., 2022), *Rspo1* (Zhang et al., 2021), *Esr1* (Li et al., 2022b), and *Sox8* (Yang et al., 2023). Accordingly, we hypothesize that the W chromosome may contain different SDGs and regulates sex determination in *P*. *sinensis* through the star genes, which is consistent with the hypothesis of “masters change, slaves remain” (Graham et al., 2003), meaning that the SDGs may be different while the genes in the midstream or downstream of the sex-determining pathway are relatively conserved.

### 4.2 A conserved ZW sex determination system presents in Trionychidae

Among the four soft-shelled turtles we investigated, the genomic similarity was more than 90% and the W chromosome was sequenced at half the depth of the Z chromosome. More importantly, all sex-linked SNPs in the four soft-shelled turtles were localized on the W chromosome, exhibiting heterozygosity in females and purity in males. Additionally, phylogenetic and divergence time results indicated that all surveyed soft-shelled turtles shared a common ancestor and that the time of sex chromosomes formation preceded the divergence of Trionychidae species. All evidences imply that a conserved ZW-type sex determination system may exist in the Trionychidae. Our results are also consistent with previous studies. Based on the genome resequencing, multiple female sex-specific markers have been developed, which could accurately identify the genetic sex of *P*. *sinensis* populations (Liang *et al*., 2019). A female-specific signal was detected on a microchromosome by comparative genomic hybridization, revealing a ZZ/ZW system presented in *A. spinifera* (Badenhorst *et al*., 2013). Quantification of gene copy numbers in the male and female genomes by qPCR assay confirmed that nine species from four genera had homologous Z chromosomes (Rovatsos *et al*., 2017). Furthermore, a female-specific sex marker developed in *A. spinifera* allowed for cross-species sex identification for *P. sinensis* (Literman et al., 2017). By constructing the chromosome-level genome of *R. swinhoei* and *P. cantorii*, the clues that they have a ZW sex-determination system have also been discovered (Liu *et al*., 2023; Ren *et al*., 2022). Collectively, these species encompass 9 of the 13 genera in Trionychidae (Shine, 2013); therefore, we have sufficient evidences and confidences to believe that there is a conserved ZW-type sex determination system in Trionychidae species. On the other hand, the high degree of consistency of the sex chromosomes of these four soft-shelled turtles suggests that they may have originated common ancestral autosomal pairs and have similar sex-determining pathways. However, the chromosome-level genome assembly of other investigated soft-shelled turtles is still needed in the future to clarify these scientific issues.

### 4.3 The origin and evolution of the sex chromosomes in Trionychidae

Studies of extant turtles revealed that most turtles have an ESD system, as if ESD seemed to be the main theme in the long evolution of turtles, with GSD only evolving independently in individual branches (Bista and Valenzuela, 2020; Modi and Crews, 2005). ESD is probably the ancestral state of amniotes (including turtles), so this transition from ESD to GSD is significant as the results of adaptive evolution of species in response to extreme environmental changes resulted in disproportionate sex ratios of offspring, thus saving species from extinction (Johnson Pokorná and Kratochvíl, 2016). Our results also demonstrated that Trionychidae species are also a specific group of turtles that transformed from ESD to GSD. The six investigated species of Trionychidae gained sex chromosomes in their common ancestor approximately 56 million years ago, earlier than the divergence time among them and later than *C. insculpta*. This well explained the possibility that *C. insculpta* has a TSD system (Webb *et al*., 1986) and is roughly consistent with the extrapolation of Rovatsos et al. that the origin of the sex chromosomes in trionychids between the split of Trionychidae and Carettochelyidae, and the basal splitting of the recent trionychids (Rovatsos *et al*., 2017).

Intriguingly, although the evolution of the main morphological features of the Trionychidae species was completed ∼120 million years ago (Li *et al*., 2015), the form of the sex chromosomes seems to be a fortuitous event, which perfectly coincides with the occurrence of the Paleocene-Eocene thermal maximum, PETM, event (Li et al., 2022a; Westerhold et al., 2020). The PETM, a major greenhouse event lasting ∼200 thousand years (Zeebe and Lourens, 2019), has led to a global temperature rise of about 5-9 °C (Glikson, 2016; Wright and Schaller, 2013). Therefore, we hypothesize that the ancestors of Trionychidae species were temperature-sensitive ESD turtles that gradually evolved into GSD populations due to the PETM event, thereby antagonizing the population sex ratio imbalance caused by the rapid increases in global temperatures. Surveys of the distribution range of extant turtles also provide some clues. The majority of Trionychidae species are found around the Tropic of Cancer and the equator. Instead, hard-shelled turtles have a broader range and appear around 60 degrees north and south latitudes (Shine, 2013). The conjecture is also consistent with a recent study of the scallop genome, which concluded that homomorphic sex chromosomes are the “normal phenomenon” in sex chromosome evolution, while heteromorphic sex chromosomes require extra evolutionary dynamics for their formation (Han et al., 2022). Altogether, the origin of the sex chromosomes of Trionychidae species is a fine example of the adaptive evolution occurred in response to extreme temperature changes and provides a reference for the studies of the species evolution of turtles, especially under the background of global warming (Jensen et al., 2018; Pike, 2014).

Alternatively, except for the maternal effects that allow TSD turtles to circumvent the imbalances of offspring sex ratio caused by the temperature variation, such as females can choose the appropriate times and places for nesting and laying eggs to avoid the effects of extreme temperatures (Du et al., 2023; Mitchell and Janzen, 2010), the embryonic self-regulation in eggs is also a critical mean (While and Wapstra, 2019; Ye et al., 2019). Briefly, the embryos can move within the eggs to select the optimal ambient temperature, thus influencing their own sex fates and maintaining the fitness sex ratios of the population. Most turtles have oval-shaped eggs (Ye *et al*., 2019; Zhu et al., 2006), while the eggs of Trionychidae species are small and rounded (Li et al., 2020; Zhu et al., 2023), which may have resulted in limited space for the embryos to move, as well as reduced thermal heterogeneity within the eggs (Cordero et al., 2018; Telemeco et al., 2016), resulting in the loss of the embryo’s ability to self-regulate during the evolutionary process. This also may be one of the driving forces for sex chromosomes formation in Trionychidae species. This may also explain why most turtles maintained the TSD system in response to dramatic changes of external temperatures throughout their long evolutionary history instead of evolving sex chromosomes.

Simultaneously, when we used *P. sinensis* as a model to study the characteristics of sex chromosomes evolution, we found that the sex chromosome evolution of *P. sinensis* conformed to the general pattern of sex chromosomes formation (Bachtrog, 2013; Zhou *et al*., 2014). The sex chromosomes of *P. sinensis* originated from a conserved chromosome pair in the ancestral TSD turtles. However, the sex-determining genes *Sry* (Koopman et al., 1991; Sinclair et al., 1990), *Dmrt1* (Smith *et al*., 2009; Yoshimoto *et al*., 2008), *Amh* (Pan *et al*., 2019; Schoch and Sues, 2015), and their homologs were not found in this conserved autosomal pair or in the sex chromosomes of *P. sinensis*. The sex chromosomes of *P. sinensis* were formed approximately ∼56 million years ago, and are the highly differentiated ancient sex chromosomes. The Z chromosome of P. sinensis retained most of the functional genes compared to the ancestral sex chromosome in TSD turtles, but two obvious inversions were also detected at both ends of the chromosome. Meanwhile, the dominant features of the W chromosome are marked degradation in contrast to the Z chromosome, with the W chromosome only having about 12 Mb roughly one-third of the Z chromosome, and a significant accumulation of DNA transposable element (LINE), indicating an extensive recombination suppression between the sex chromosomes of *P. sinensis*.

Recombination suppression is thought to occur after the fixation of sex-determining genes and is an indispensable driving force in the evolution of sex chromosomes (Charlesworth, 2017; Rifkin et al., 2021). The chromosomal inversion and transposon accumulation are thought to induce the development of recombination suppression (Charlesworth *et al*., 2005). Following the initial recombination, the chromosomal inversion decreases the recombination rate of sex-determining genes and further expands the nonrecombining regions in the sex chromosomes (Furman et al., 2020; Otto et al., 2011). Previous studies also have shown that massive transposable elements (TEs) are found at the border between recombinant and nonrecombinant regions of both mammalian and avian sex chromosomes (Iwase et al., 2003; Xu et al., 2019a). The insertion of TEs near the sex-determining locus can utilize the divergences between sex chromosomes to suppress recombination (Kent et al., 2017). Besides, our findings also revealed that the W chromosome of *P. sinensis* has a relatively high rate of non-synonymous substitution to the Z chromosome (or ancestral sex chromosome in TSD turtles) and autosomes. Hill-Robertson interference argues that recombination suppression leads to the accumulation of deleterious mutations and a decrease in adaptive levels (Felsenstein, 1974), thereby resulting in the loss of functional genes and the degeneration of sex chromosomes. Overall, recombination suppression benefits sex-determining genes and sex-linked genes that are inherited in one sex, allowing sex chromosomes to evolve toward heterochromatin (Ponnikas et al., 2018; Rice, 1987).

### 4.4 Single origin of the Trionychidae species

Based on the above findings, the sex chromosomes of the six investigated softshell turtles have a single origin in a common ancestor predating the species divergence times among them, and the high identity among the sex chromosome fragments of the four investigated softshell turtles suggesting their sex chromosomes probably originated from the same autosomal pairs in the common ancestor, all support the view that the soft-shelled turtles acquired their sex chromosomes from a common ancestor before dispersing around the world. These clues raise our great interests in the origin and radiation of Trionychidae species. We believe that the ancestor of the extant Trionychidae species has a high probability of originating in Southeast Asia. Extant Trionychidae species have a relatively broad distribution across the tropical to warm temperate portions of Asia, North America, and Africa (Georgalis and Joyce, 2017; Vitek and Joyce, 2015). However, we first rule out the possibility of an African origin. Because Africa bordered Eurasia very late around five million years ago, and time tree results showed that *P. cantorii*, *R. swinhoei*, *P. sinensis*, and *P. steindachneri* had already diverged at this time, which contradicts previous inferences of a single common ancestor for all extant soft-shelled turtles. Moreover, it would not be North America, since the ancestor of Trionychidae species was first separated from that of *C. insculpta*, and the first divergent species in Trionychidae was *P. cantorii*, both of which are primarily distributed in Southeast Asia (Shine, 2013). Fossil studies also showed that the ancestors of *C. insculpta* crossed the Wallace Line from Southeast Asia into areas of New Guinea and Australia no later than the late Miocene (Joyce, 2014; Rule et al., 2022).

We think that the common ancestor of extant soft-shelled turtles originated in Southeast Asia, which is strongly consistent with historical climate changes, continental plate movements and the current molecular evidences. Due to the global PETM event about 56 million years ago (Glikson, 2016; Wright and Schaller, 2013), temperatures near the equator rose rapidly, causing the common ancestor of the extant soft-shelled turtles in Southeast Asia to gradually evolve sex chromosomes and disperse to higher latitudes to avoid possible population extinction as a result of the high-temperature event. PETM events are recognized as an important driver of the radial evolution of species (Bowen et al., 2002; Gingerich, 2006), such as the smaller body size of some mammals (Secord et al., 2012) and the evolution of terrestrial artiodactyls to cetaceans (Thewissen et al., 2007; Yuan et al., 2021). Diversity and dispersal of soft-shelled turtles significantly accelerated during the Paleocene global warming period through a study of mitochondrial genes (Le et al., 2014). Subsequently, this sub-group spread from Eurasia to North America via the Bering Land Bridge and gradually evolved into the species of the genus *Apalone*. During the Paleocene and Eocene, tropical flora widely distributed in the Northern Hemisphere, and the Bering Land Bridge became a major conduit for species dispersal between Eurasia and the North American, including 10 reptile dispersal events (Jiang et al., 2019). Meanwhile, another group of soft-shell turtle ancestors spread and evolved during the gradual approach and eventual collision of the Indian subcontinent with the Eurasian in the Eocene (Ali and Aitchison, 2008; van Hinsbergen et al., 2012). Similar results were confirmed in a fossil study (Smith et al., 2016). Biological dispersal between the Indian subcontinent and mainland Asian continent was a dynamic process that accelerated at 44 Ma, peaked in the mid-Miocene, and declined after 14 Ma (Klaus et al., 2016). Finally, when the African bordered the Eurasian, Trionychidae species from regions near India subcontinent migrated to Africa and gradually diverged. The reason for this is that the genome and mitochondrial phylogenetic tree results indicated that *A. spinifera* and *A. ferox* are closely related to *R. swinhoei* and *P. sinensis* distributed in Asia, whereas *Trionyx triunguis* is more closely related to *C. indica* and *P. cantorii* presented in India. Taken together, our results support that the common ancestor of extant soft-shelled turtles disperses to higher latitudes centered on the Southeast Asian driven by the PETM event, and subsequently enters North America via the Bering land bridge. However, another group moves into the Indian subcontinent and then spreads into Africa (Figure 7).

**Figure 7.**
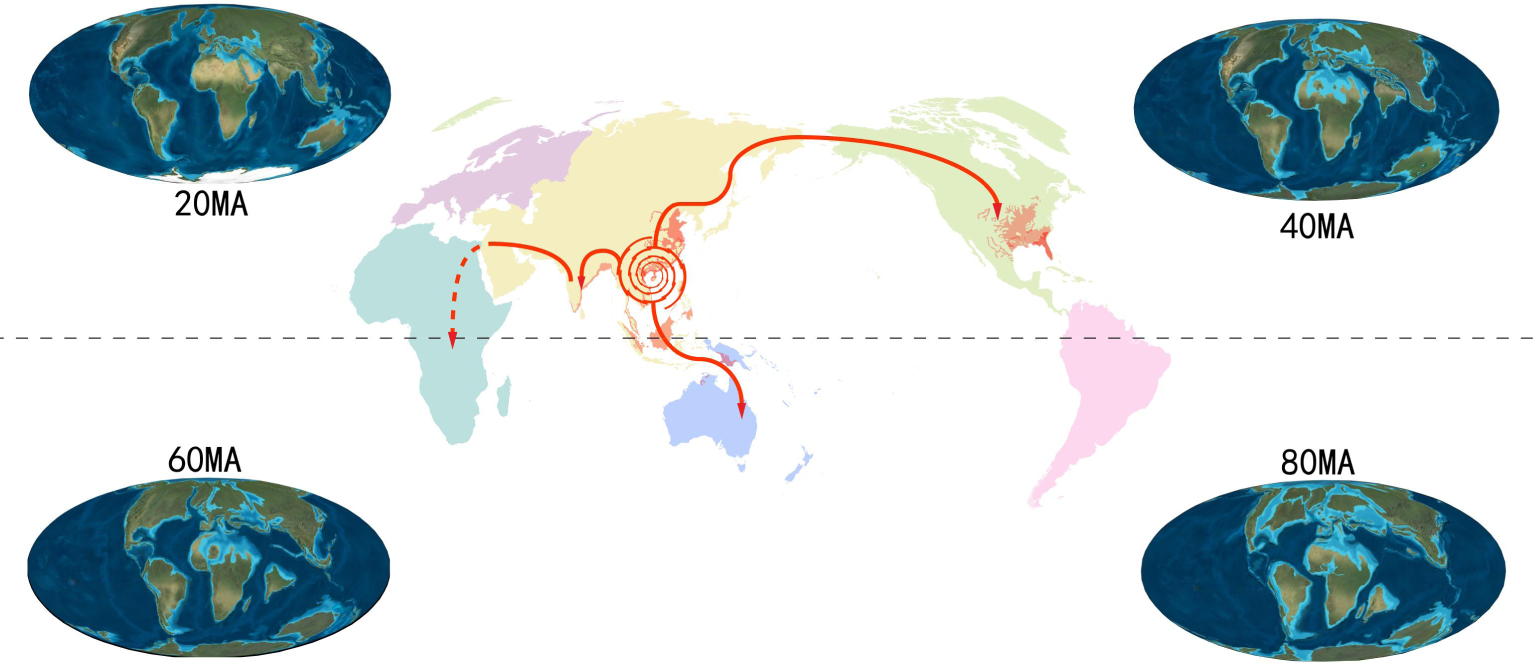
Evolutionary pathways of extant soft-shelled turtles.

## 5. Conclusion

In summary, we assembled the first chromosome-level genome and the first sex chromosomes of *P. sinensis*, and rejected the involvement of *Dmrt1*’s SDGs or dosage compensation effect in the sex determination of *P. sinensis* ZW system. Moreover, comparative genomic studies reveal a conserved ZW sex-determination system presented in Trionychidae species, as well as the single origin of the sex chromosomes and species of Trionychidae species.

